# Reciprocal macrophage-MSC crosstalk drives immunomodulatory and regenerative phenotypes in a mineralized collagen scaffold

**DOI:** 10.64898/2026.03.10.710803

**Authors:** Vasiliki Kolliopoulos, Maxwell Polanek, Hashni E. Vidana Gamage, Melisande Wong Yan Ling, Aleczandria Tiffany, Erik R. Nelson, Kara L. Spiller, Brendan A.C. Harley

## Abstract

Critical sized craniomaxillofacial bone defects do not heal naturally and often exhibit chronic inflammatory responses that restrict regeneration. It is increasingly apparent that biomaterials must facilitate dynamic crosstalk between immune cells, such as macrophages, and osteoprogenitors to resolve inflammation and accelerate regeneration. Here, we evaluate interactions between macrophages in a neutral (M0) or pro-inflammatory (M1) state with mesenchymal stem cells (MSCs) in a basal or licensed state within a mineralized collagen scaffold. We reveal that MSC-macrophage crosstalk influences significant changes in osteoprogenitor cell differentiation and immune cell polarization. Notably, crosstalk between MSCs and macrophages drives an early-stage inflammatory response, which enhances the immunomodulatory activity of MSCs via secretion of IL-6, an effect that is heightened for already licensed MSCs. The presence of macrophages in the co-cultures upregulated osteogenic (*ALPL*, *BMP2*, *COL1A2*, and *RUNX2*) and angiogenic genes (*ANGPT1*) in basal MSC groups. Further, MSC-macrophage interactions subsequently drive increased M2-like macrophage polarization as early as 7 days of culture, as indicated by surface marker expression. These findings show that biomaterial scaffolds can be leveraged as mediators of MSC-mediated immunomodulation with an emphasis on achieving early-stage pro-inflammatory phenotypes that drive subsequent macrophage polarization and markers of increased regenerative potency.

## 1. Introduction

Critical sized craniomaxillofacial (CMF) bone defects can be large in size (>25 mm) and irregular in shape, making them challenging to heal. They arise most commonly from congenital abnormalities, trauma, or tumor resection[1, 2]. Current standards of repair include traditional grafting techniques such as autografts and allografts[3, 4]. However, these suffer from challenges such as donor-site morbidity, limited availability for critical sized defects, the need for multiple surgeries, and risk of infection. To overcome these limitations, tissue engineering efforts have focused on development of biomaterials that recruit endogenous cells – including stem cells – to potentiate bone regeneration[5–7]. However, the initial inflammatory status immediately after injury, as well as the reciprocal crosstalk between inflammatory and regenerative cells within a biomaterial implant, may strongly influence regenerative potential.

Tissue repair often follows three phases: inflammation, tissue formation, and remodeling[8, 9]. Regulation of the inflammatory phase has gained significant interest as this phase is important for inhibiting infection, clearing necrotic tissue, and influencing downstream tissue formation and remodeling[10, 11]. Macrophages play crucial roles in each phase of healing, accumulating in the damaged tissue as they are recruited by inflammatory stimuli and exhibiting tremendous plasticity in response to surrounding environments. Under normal healing conditions, macrophages first adopt a pro-inflammatory “M1” phenotype clearing damaged tissue, secreting pro-inflammatory markers such as tumor necrosis factor-α (TNF-α) and interleukin-6 (IL-6) and initiating the healing process through the stimulation of angiogenesis. These M1 macrophages can then polarize into a diverse population of macrophages collectively referred to as “M2” which stabilize angiogenesis and orchestrate extracellular matrix assembly[12], although the extent of the diversity of this population is not known. When a biomaterial is implanted into a wound site, it is exposed to an inflammatory environment driven by M1 macrophages and their inflammatory secretome. A regenerative biomaterial may have to recruit, retain, and support the pro-regenerative activity of osteoprogenitors in the context of extended inflammatory conditions. Therefore, there is an opportunity to consider biomaterial solutions that can ameliorate a macrophage-driven inflammatory state and facilitate osteoprogenitor activity to accelerate bone regeneration.

In addition to their potential as osteoprogenitors for craniofacial bone regeneration applications, human mesenchymal stem cells (hMSCs) are responsive to their local inflammatory environment. MSCs can take on a “licensed” phenotype when exposed to inflammatory factors such as interferon-γ (IFN-γ) and TNF-α. Licensed MSCs can differentially modulate their surrounding microenvironment, including immune cells such as macrophages, T-cells, and B-cells, via the secretion of paracrine factors such as prostaglandin E_2_ (PGE_2_), interleukin-6 (IL-6), and indoleamine 2,3-dioxygenase (IDO1)[13–17]. Not surprisingly, macrophages can also promote changes in hMSC osteogenesis via the COX2-PGE_2_ pathway[18]. Although numerous studies have focused on the effects of MSC licensing, reciprocal crosstalk between clinically relevant macrophages and osteoprogenitors remains to be investigated in a bone-mimicking microenvironment.

We have previously described a mineralized collagen biomaterial for critical-sized CMF bone regeneration that promotes MSC osteogenic differentiation *in vitro* and induces mineral and bone formation *in vivo* without the use of exogenously added osteogenic factors[19]. We showed scaffold proteoglycan composition and pore anisotropy each shape MSC osteogenic and immunomodulatory phenotype. While heparin-modified scaffolds maximally promoted osteogenic activity, scaffolds containing chondroitin-6-sulfate promoted production of a pro-angiogenic and immunomodulatory phenotype in licensed hMSCs[20, 21]. Interestingly, we also observed that anisotropic (aligned) scaffold pore architecture elevated immunomodulatory potential under licensed conditions[20, 22]. These findings suggest that while licensing is a driving factor of MSC immunomodulatory potential, MSC activity can be further tuned via material structure and composition. Separately, we used the mineralized collagen scaffold to evaluate indirect (conditioned media) vs. direct co-culture of human THP-1 monocytic cell line-derived macrophages and MSCs[22]. We observed upregulated MSC immunomodulatory activity in the presence of conditioned media from macrophages as well as an elevated osteogenic response to M1-associated paracrine factors. However, in direct THP-1/MSC co-cultures, we saw a reduced upregulation of immunomodulatory and osteogenic factors but significantly upregulated matrix remodeling genes in MSCs. This well-characterized scaffold environment provides a 3D *in vitro* platform to interrogate the role of scaffold architecture on MSC-macrophage crosstalk interactions that occur in the context of inflammation and regeneration[21–23]. Finally, we have also shown that the prior polarization state of macrophages alters their ability to respond to regenerative cues, in that pro-inflammatory M1 macrophages were more sensitive to the reparative cytokine interleukin-4 (IL-4) and switched to a distinct M2-like phenotype with potentially enhanced regenerative capacity[24]. There is a need to better understand how macrophage and MSC phenotypes influence their interactions within relevant 3D environments.

Herein, we report the effect of direct (co-culture) interactions between human MSCs and primary human monocyte-derived macrophages within a mineralized collagen scaffold *in vitro*. We describe the consequences of MSC-macrophage crosstalk as a function of both initial macrophage phenotype (non-activated M0 compared to the pro-inflammatory M1) as well as initial MSC inflammatory state (basal compared to licensed). We hypothesize that an initial inflammatory macrophage phenotype or primed MSC state will enhance MSC-mediated immunomodulation and macrophage pro-healing polarization, respectively. This study provides new insight regarding the driving force of MSC immunomodulatory activity as well as shifts in macrophage inflammatory phenotype within mineralized collagen scaffolds as a result of MSC-macrophage crosstalk. These insights will help inform improved design rules for regenerative biomaterials that support *in vivo* craniofacial bone regeneration.

## 2. Materials and Methods

### 2.1. Experimental design

This study used direct co-culture to investigate interactions between basal or licensed human MSCs with non-activated (M0) or pro-inflammatory (M1) activated primary human macrophages (**Figure 1**). The secretion of osteogenic and immunomodulatory factors as well as MSC gene expression profiles within the mineralized collagen scaffolds were assessed as a function of initial MSC inflammatory status and exposure to M0 or M1 macrophages. We characterized dynamic changes in macrophage phenotype in response to hMSCs via surface marker expression and gene expression.

**Figure 1:**
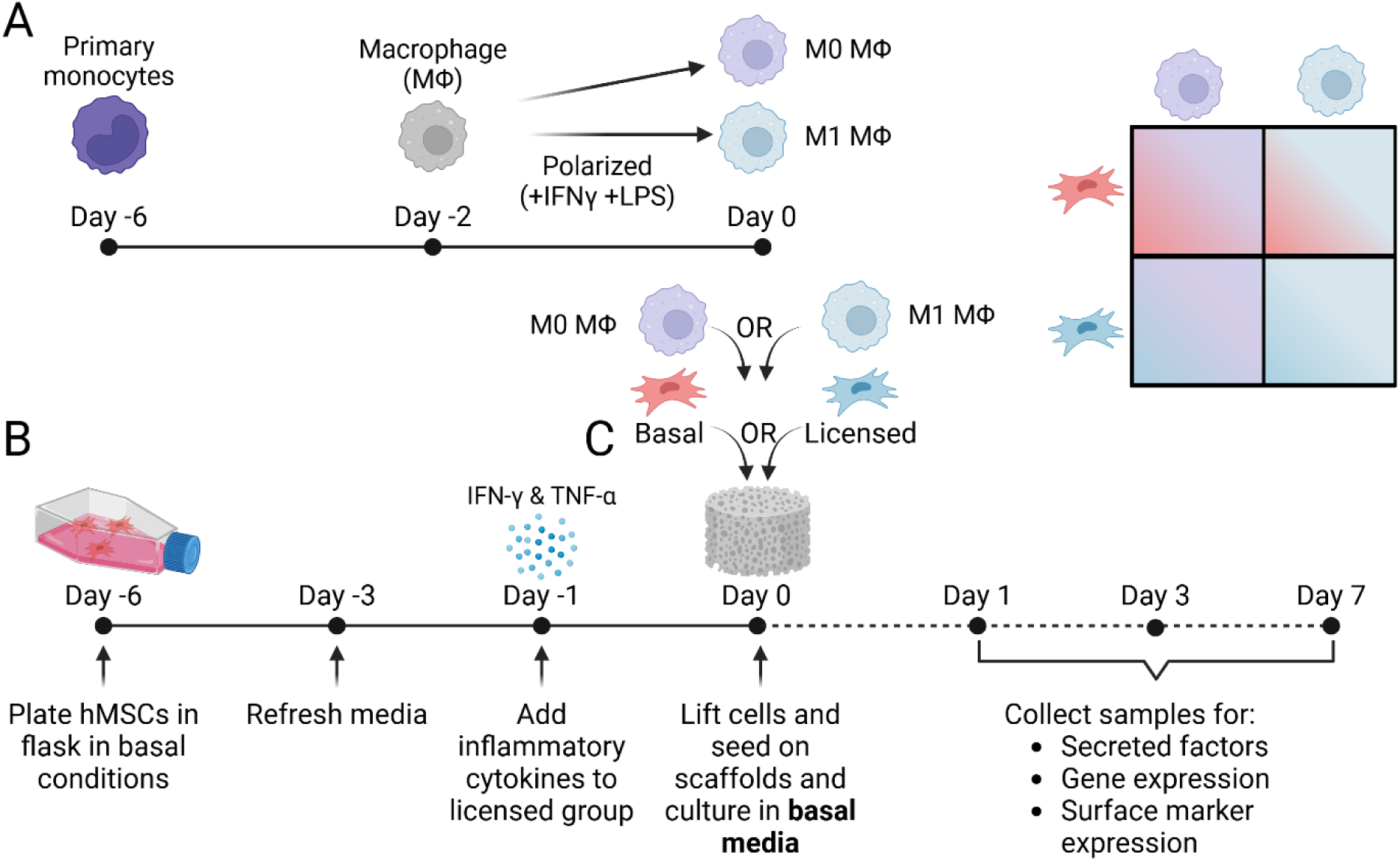
(A) Human peripheral blood monocytes were cultured, differentiated into macrophages, and polarized to non-activated (M0) or pro-inflammatory (M1) macrophages. (B) Human mesenchymal stem cells (hMSCs) were cultured in basal media or stimulated with inflammatory cytokines 24 hours prior to seeding on mineralized collagen scaffolds. (C) Mineralized collagen scaffolds seeded with basal or licensed hMSCs and with M0 or M1 macrophages were maintained *in vitro* for 7 days to define the influence of hMSC-macrophage co-culture on hMSC osteogenic response and macrophage polarization.

### 2.2. Fabrication of mineralized collagen-glycosaminoglycan scaffolds with aligned pores

Mineralized collagen-glycosaminoglycan scaffolds with aligned pores were fabricated from a mineralized collagen precursor suspension as previously described[21, 23, 25, 26]. Briefly, type I bovine collagen (1.9 w/v% Collagen Matrix Inc., New Jersey USA) suspended in a mineral buffer solution (0.1456 M phosphoric acid/0.037 M calcium hydroxide) was homogenized and allowed to hydrate at 4°C overnight. The collagen suspension was then homogenized with calcium salts (calcium hydroxide and calcium nitrate tetrahydrate, Sigma-Aldrich), and chondroitin-6-sulfate (CS6) (0.84 w/v%, Chondroitin sulfate sodium salt, CAS Number 9082-07-9, Spectrum Chemicals). The mineralized collagen suspension was then transferred to a Teflon mold with a copper base and lyophilized into porous scaffolds with aligned pores using a Genesis freeze-dryer (VirTis, Gardener, New York USA). The suspensions were cooled at a constant rate of 1 °C/min from 20°C to −10°C, then held at −10°C for 2 hours, followed by sublimation of the ice crystals at 0°C and 0.2 Torr, resulting in a porous scaffold network. The porous scaffolds were then trimmed to generate cylindrical scaffolds with 6 mm by 1.5 mm in height for the direct co-culture studies.

### 2.3. Sterilization, hydration, and scaffold crosslinking

Scaffolds were sterilized via ethylene oxide treatment for 12 hours using a AN74i Anprolene gas sterilizer (Andersen Sterilizers Inc., Haw River, North Carolina USA). All subsequent steps were conducted in aseptic environments. Sterile scaffolds were hydrated and crosslinked as previously described[21, 23, 27, 28]. First scaffolds were placed in 100% sterile ethanol for 2 hours and subsequently washed in PBS for 1 hour on a shaker at room temperature. The scaffolds were then crosslinked using EDC-NHS chemistry for 1.5 hour while shaking at room temperature. Following crosslinking the scaffolds were thoroughly washed in PBS for 1 hour and finally soaked in basal growth media for 48 hours prior to cell seeding with a media change after 24 hours.

### 2.4. Cell culture, priming, and seeding of human mesenchymal stem cells and differentiating and polarizing macrophages for indirect co-cultures

#### 2.4.1. Human mesenchymal stem cell culture and priming

Human bone marrow derived mesenchymal stem cells (hMSCs) of a 20-year-old African American female (RoosterBio, Frederick, MD, USA) were expanded from passage 4 to passage 5 using RoosterNurish expansion media for the first 3 days followed by 3 more days of culture media containing low glucose DMEM and glutamine, 10% mesenchymal stem cell fetal bovine serum (Gemini, California, USA), and 1% antibiotic-antimycotic (Gibco, Massachusetts, USA) in an incubator at 37°C and 5% CO_2_. On day 5 of culture hMSCs were licensed with 20 ng/mL IFN-γ and 10 ng/mL TNF-α (cyt-206 and cyt-223 respectively, ProSpec Protein Specialists) for 24 hours prior to seeding onto the scaffolds[13, 20]. Basal hMSCs were generated by culturing in basal media through day 6 of cell expansion prior to seeding on the scaffolds.

### 2.5. Differentiation and culture of primary human peripheral blood monocytes with human mesenchymal stem cells in direct scaffold co-culture

#### 2.5.1. Peripheral blood human monocyte culture and differentiation to macrophages

Human primary peripheral blood monocytes (70034, STEMCELL Technologies, Vancouver, Canada) were cultured, differentiated, and activated into non-activated M0 macrophages as per the manufacturer’s instructions. Briefly, monocytes were placed in non-tissue culture treated T175 flasks at a concentration of 1 million cells/mL and differentiated to macrophages in ImmunoCult™-SF Macrophage Medium (10961, STEMCELL Technologies, Vancouver, Canada) supplemented with 50 ng/mL recombinant human M-CSF (78057, STEMCELL Technologies, Vancouver, Canada) for 4 days at 37°C and 5% CO_2_ in an incubator. Following 4 days of differentiation, half of the original medium containing M-CSF was added and the cells were cultured for an additional 2 days prior to seeding with no additional cytokines to generate non-activated (M0) macrophages or supplemented with 50 ng/mL IFN-γ and 10 ng/mL of LPS to generate pro-inflammatory activated (M1) macrophages. Following a total of 6 days of culture in the flasks, macrophages were lifted using StemPro Accutase (ThermoFisher Scientific) for 15 minutes at 37°C and 5% CO_2_ in an incubator. The Accutase was neutralized with RPMI medium and cells were gently scraped. Cells were then centrifuged at 300 g for 10 minutes. Cells were then resuspended in fresh RPMI medium supplemented with 50 ng/mL recombinant human M-CSF.

#### 2.5.2. Seeding of hMSCs and macrophages on mineralized collagen scaffolds

hMSCs and macrophages were co-cultured within mineralized collagen scaffolds at a 1:3 ratio for 7 days. Basal or licensed hMSCs were resuspended to a final concentration of 10^7^ cells/mL while M0 or M1 macrophages at a final concentration of 3×10^7^ cells/mL. Each scaffold was placed in a separate well of a 24-well plate and seeded with 5 μL of hMSCs and 5 μL of macrophages resulting in final seeding densities of 50,000 hMSCs and 150,000 macrophages per scaffold. Cell seeded scaffolds were incubated at 37 °C and 5% CO2 for 1 hour. The following cell groups were studied: basal hMSCs + M0 macrophages (M0B); licensed hMSCs + M0 macrophages (M0L); basal hMSCs + M1 macrophages (M1B); licensed hMSCs + M1 macrophages (M1L). All co-culture groups were maintained in 50% hMSC basal media and 50% RPMI media supplemented with MCSF. Control groups included single culture scaffolds with only basal or licensed hMSCs or M0 or M1 macrophages at their respective seeding densities. Control groups containing hMSCs were cultured in basal hMSC media and control groups containing macrophages were cultured in RPMI media supplemented with MCSF. Media was changed every 3 days.

### 2.6. Flow cytometry

At days 1, 3, and 7 of direct co-culture of hMSCs and macrophages, cells were removed from the scaffolds for macrophage phenotypic characterization via flow cytometry. Briefly, scaffolds were washed in PBS and then quartered. Scaffolds were then soaked in TrypLE for 5 minutes 37°C and 5% CO_2_ in an incubator. Subsequently, an equal volume of 5 mg/mL collagenase was added to each scaffold and incubated for 20 minutes at 37°C and 5% CO_2_ on a shaker. The TrypLE and collagenase was neutralized with RPMI media and the solution was filtered through FACS tubes with a 35-μm strainer cap at 1100 rpm for 5 minutes. The solution was aspirated with a mixed population of isolated hMSCs and macrophages resuspended in PBS prior to staining.

A fixable live/dead marker was used to stain cells for viability according to manufacturer’s instructions (Invitrogen) in PBS for 20 minutes at 4°C in the dark. Cells were then washed for 5 minutes, at 500 g at 4°C with FACS buffer (PBS containing 2% Fetal Bovine Serum and 1% Penicillin-Streptomycin). Cells were incubated with human Fc blocker in FACS buffer for 10 minutes at 4°C and subsequently incubated with flow cytometry antibody specific for human CD80 (B7-1) and CD206 (19.2) in FACS buffer for 20 minutes at 4°C in the dark. Antibodies were used at a working dilution of 1:100. The antibody panel are listed in **Supp. Table 3** After incubation cells were washed three times with FACS buffer and fixed with 4% formalin. Flow cytometry was performed on BD LSRII (BD Biosciences) and BD FACSymphony A1 (BD Biosciences). The data were analyzed by FCS Express software. Representative gating strategy is shown in **Supp. Figure 1**.

### 2.7. Secretome quantification from hMSC and macrophage direct co-culture

Soluble osteoprotegerin (OPG) and immunomodulatory factors IDO1, PGE_2_, IL-6, and CCL22 were quantified via their respective ELISAs (OPG – DY805, IL6 – DY206, IDO - DY6030, PGE_2_ – KGE004B, CCL22 - DY336, R&D Systems, Minneapolis, MN, USA) (**Figure 5**). Media was collected every 3 days throughout the 7-day culture period and pooled (day 1; day 3; days 6-7) to generate a cumulative release profile normalized to a blank media control (n=3).

### 2.8. RNA extraction and NanoString gene expression analysis of hMSC and macrophages in direct co-culture

Scaffolds seeded with hMSCs and macrophages were used to examine gene expression as previously described (n=3)[20, 29]. Each scaffold was quartered and placed into phasemaker™ tube (ThermoFisher Scientific, Massachusetts, USA) with 1 mL of TRIzol™ Reagent (ThermoFisher Scientific, Massachusetts, USA) and vortexed. To the scaffold TRizol solution, 200 μL of chloroform was added, vortexed and allowed to rest for 5 minutes. Samples were centrifuged for 15 minutes at 4°C and the supernatant containing the RNA was collected and diluted by half in 70% ethanol. The entire suspension was transferred to RNA extraction columns of the RNA Clean & Concentrator-5 kit (Zymo Research). All subsequent steps were followed per manufacturer’s instructions. RNA concentration and purity was measured using a NanoDrop spectrophotometer where all 260/280 nm ratios were all above 1.8 and below 2.3. Samples were assessed with two custom NanoString panels. The first custom panel of 38 mRNA probes was used to identify the osteogenic, immunomodulatory, and angiogenic response by hMSCs (**Supp. Table 1**). The second panel consisted of 72 genes fully characterizing macrophage phenotype specific gene quantification allowing for macrophage phenotyping **(Supp. Table 2**), as previously described[30]. Raw data from both panels were normalized to positive and negative internal controls and then to two housekeeping genes (*GAPDH* and *TBP*). This data was further normalized to a day 0 control (n=3) and expression levels are depicted as a log base 2-fold change. Although gene expression cannot be reliably isolated to one cell type in the co-cultures, further analysis of the gene data set was conducted to isolate MSC-only and MΦ-only genes. Cell-specific genes were identified by taking the difference of the experimental sample with the individual cell controls. Genes that were negative for the one cell type were considered cell-specific for the other. Further down selection of cell-specific genes was conducted by normalizing the log2 values of the experimental samples to the day 0 single culture control and propagating the error (Log2(exp)-log2(Day0-single culture)). If the sum of the individual controls was greater than one standard deviation from the experimental samples these genes were selected as genes of interest.

### 2.9. Statistics

Statistical analysis was performed with RStudio (RStudio, Massachusetts, USA) and OriginPro (OriginPro, Massachusetts, USA) software packages. A p-value less than 0.05 was considered statistically significant. Statistics were executed per time point. No outliers were removed. All plots were created in OriginPro. Error bars for each group were represented as mean ± standard deviation. For comparison between co-cultures (M0B, M1B, M0L, and M1L) at each time point, the assumptions of normality and equal variances were evaluated using the Shapiro-Wilk normality test and Levene’s test, respectively. For comparisons to single culture controls at each time point (only shown in **Supp. Figure 2 and 3**), a similar evaluation was performed. If the dataset (including single culture controls) was normal and had equal variances, a two-sample T-test was performed to evaluate differences from the single culture controls. Otherwise, a Mann-Whitney test was performed. If the dataset was both normal and had equal variances, a two-way ANOVA with Tukey post-hoc was performed in Origin to assess significance using MSC treatment and macrophage polarization as the independent variables. ELISA protein assays were performed with n=4 and were normalized to appropriate media culture controls. Nanostring gene expression assays were performed with n=3 and were normalized to either a macrophage or MSC single culture control.

**Figure 2:**
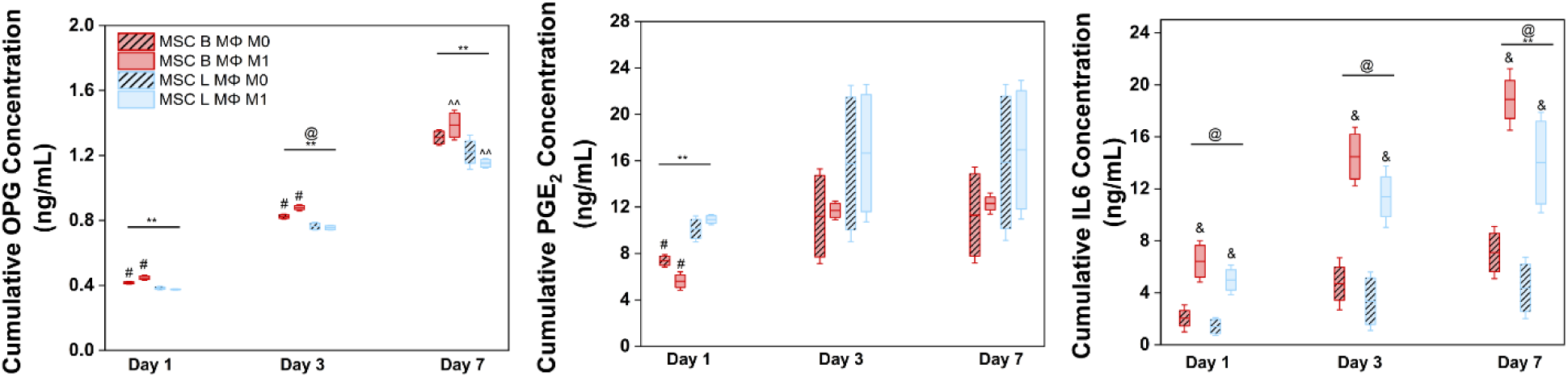
ELISAs. Mineralized collagen scaffolds seeded with basal or licensed hMSCs were co-cultured with M0 or M1 primary macrophages. Media was collected every 3 days and analyzed via ELISAs. Osteoprotegerin (OPG), an osteoclast inhibitor; prostaglandin-E_2_ (PGE_2_), a key immunomodulatory cytokine acting on macrophages; and interleukin-6 (IL6), another key immunostimulatory cytokine, were quantified as a function of MSC treatment and macrophage phenotype. ** denotes overall significance (p < 0.05) between MSC treatments at the indicated timepoint, @ denotes overall significance (p < 0.05) between macrophage polarization status at the indicated timepoint, ^^ denotes significance (p < 0.05) between indicated group and all other groups of other MSC treatment, & denotes significance (p < 0.05) between indicated group and all other groups of other macrophage polarization, and # denotes significance (p < 0.05) of indicated group and all other groups at that timepoint.

**Figure 3:**
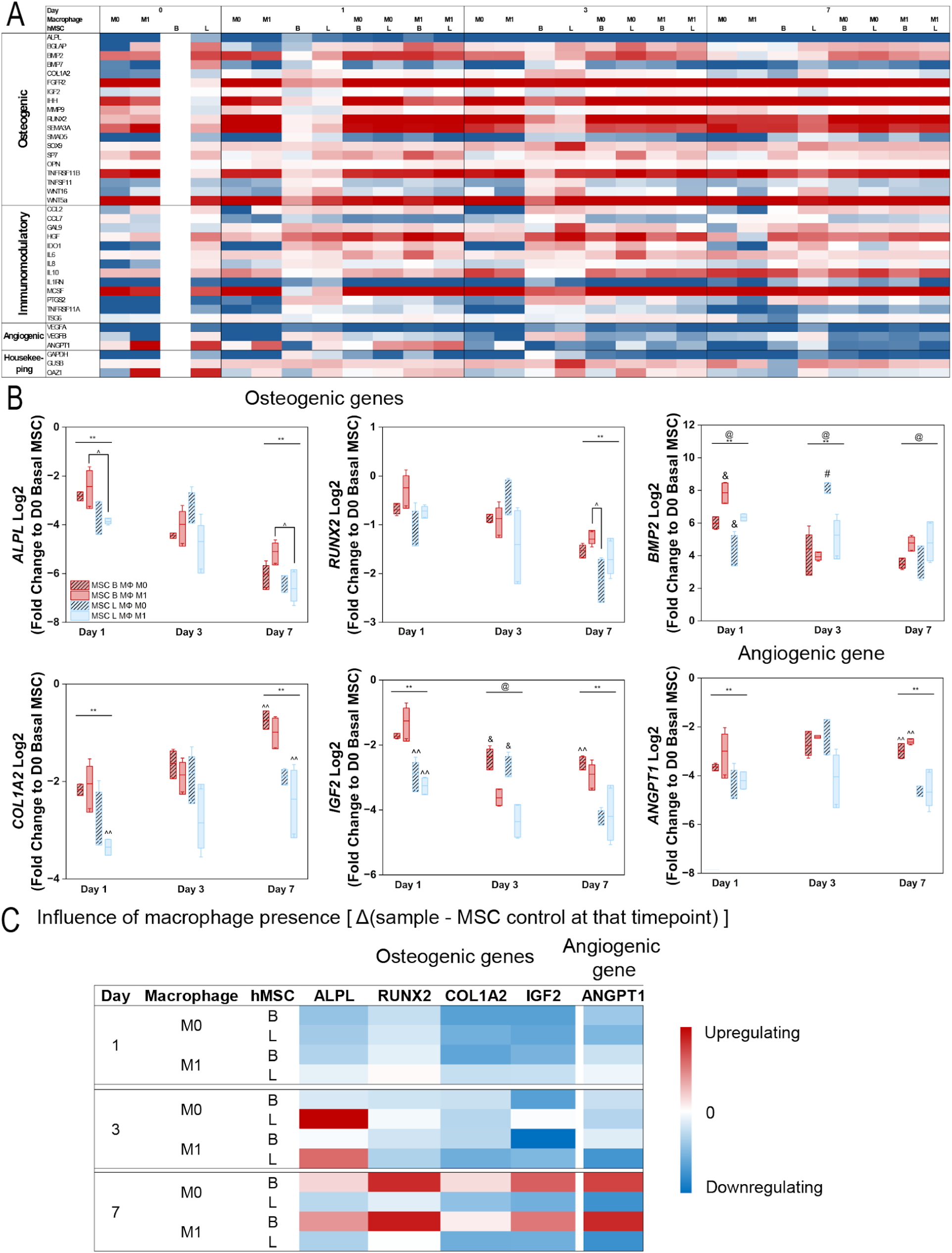
Patterns of MSC gene expression as a function of inflammatory licensing and macrophage co-culture. A custom NanoString code set was used to quantify MSC gene expression in single culture controls and as a function of MSCs basal vs. licensed status and in response to co-culture with primary macrophage (ΜΦ) phenotype (M0 vs. M1). A subset of genes was identified in the MSC panel that were identified as MSC only genes and where the co-cultures displayed > or < than 1 standard deviation from the single culture MSC controls. (A) All genes displayed as a fold change compared to basal MSC gene expression prior to seeding (n=3). (B) Co-culture gene expression patterns displayed as a fold change compared to basal MSC gene expression prior to seeding (n=3). ^ denotes significance (p < 0.05) between indicated groups of different treatment (basal or licensed), ** denotes overall significance (p < 0.05) between MSC treatments at the indicated timepoint, @ denotes overall significance (p < 0.05) between macrophage polarizations at the indicated timepoint, ^^ denotes significance (p < 0.05) between indicated group and all other groups of other MSC treatment, and & denotes significance (p < 0.05) between indicated group and all other groups of other macrophage polarization. (C) The effect of MΦs co-culture quantified by the difference of co-culture vs. MSC control at each timepoint. A red color indicates macrophage co-culture enhances MSC gene expression compared to single culture control. A blue color indicates macrophage co-culture decreases indicated MSC gene expression compared to single culture control.

## 3. Results

### 3.1 M1 macrophages enhance expression of OPG and IL-6 in MSC co-cultures

First, we assessed the effect of starting macrophage phenotype on MSC behavior in terms of secretion of osteogenic and immunomodulatory soluble factors. We previously showed MSCs in mineralized collagen scaffolds secrete the glycoprotein OPG (osteoprotegerin) to inhibit osteoclast activity and enhance regenerative potency[26, 31, 32]. While immunomodulatory factors such as PGE_2_ and IL-6, known to be produced by macrophages, play critical roles in the resolution of inflammation[33–35], these factors also influence bone formation and resorption. Specifically, PGE_2_ has been shown to enhance bone mass while IL-6 stimulated osteoclast activation via RANKL expression as well as bone formation via osteocyte activity[36, 37]. Here, OPG secretion was significantly higher for basal vs. licensed hMSC co-cultures at all time points (p < 0.05) (**Figure 2** and **Supp. Figure 2**). Interestingly, within basal hMSCs co-cultures, inclusion of M1 macrophages drove significantly (p < 0.05) higher OPG secretion at days 1 and 3 compared to M0 macrophages, as well as non-significantly increased OPG secretion at day 7. MSC licensing drove increased secretion of immunomodulatory lipid PGE_2_ than in basal co-cultures at all time points, however this was only significant (p < 0.05) on day 1. IL-6 secretion by hMSCs and macrophage co-cultures was significantly increased (p < 0.05) in co-cultures containing M1 macrophages while MSC licensing significantly (p < 0.05) decreased IL-6 secretion (compared to basal co-cultures) by day 7.

### 3.2. Initial MSC licensing status dominates downstream gene expression of MSCs

We subsequently examined the effect of MSC licensing and macrophage polarization state on gene expression across a range of processes important for tissue regeneration. While MSC licensing drove MSC gene expression, macrophage co-culture had a significant effect on gene expression. Expression levels of genes associated with osteogenesis (*ALPL*, *COL1A2*, and *IGF2*) as well as angiogenesis (*ANGPT1*) were significantly decreased in cultures containing licensed MSCs both early (day 1) and late (day 7) in the culture (p < 0.05) compared to day 0 basal expression (**Figure 3A**). Expression of osteogenic gene *RUNX2* was significantly (p < 0.05) decreased by MSC licensing compared to the basal groups on day 7. Notably, although these genes did not display significant changes as a function of macrophage phenotype (e.g. M0 vs. M1), the presence of macrophages in the co-culture upregulated osteogenic (*ALPL*, *BMP2*, *COL1A2*, and *RUNX2*) and angiogenic genes (*ANGPT1*) in basal groups compared to MSC only controls on day 7 (**Supp. Figure 3**). *IGF2* expression was significantly higher (p < 0.05) in co-cultures with M0 macrophages on day 3 regardless of MSC licensing state (**Figure 3B**). The majority of the 11 osteogenic, angiogenic, and immunomodulatory genes considered in this study were downregulated (8-10) vs. upregulated (1-3) in co-culture conditions. For upregulated genes, co-cultures containing licensed MSCs exhibited the highest average fold changes following day 3 of culture. Interestingly, when we isolate the effect of macrophages in the co-cultures, we observe that macrophages drive an early-stage osteogenic response in licensed MSC groups, while a more profound osteogenic response across multiple genes occurs in the basal groups at later stages of culture (**Figure 3C**).

### 3.3. Macrophage gene expression is shaped by initial macrophage polarization state and MSC licensing

We then examined shifts in macrophage phenotype in response to MSC co-culture. Several macrophage-associated genes showed significant increases in expression in the presence of licensed MSCs (p < 0.05) compared to the day 0 control, notably those associated with M1 macrophage polarization (*CCL8,* and *TNF* on days 1 and 7, and *IL1B* on day 7, and *CD80* across the culture period) and M2a macrophage polarization. This was also seen for genes associated with the IL4/IL13-stimulated phenotype, (*CCL18* across the culture period), proliferation and immune regulation (*BTG1* on days 3 and 7 of culture, and *STAT3* on day 3 of culture) (**Figure 4A**). The presence of licensed MSCs enhanced expression of M1-associated *BTG1, CCL8, CD80, IDO1, IL1B, STAT3*, and *TNF* on day 1, compared to day 0 controls (**Figure 4B**). While MSC licensing enhanced *BTG1* macrophage gene expression through day 3 and *IL1B* expression at day 7, co-culture with basal MSCs downregulated the expression of *CCL18* and *CD80* on days 3 and 7 compared to licensed groups. **Supplemental Figure 3** also depicts macrophage-only and MSC-only control group gene expression for the selected genes seen in **Figure 4**. This suggests that co-culture of macrophages with MSCs promoted macrophage-mediated inflammatory phenotypes with the most profound effect observed for co-cultures with licensed MSCs (**Figure 4C**).

**Figure 4:**
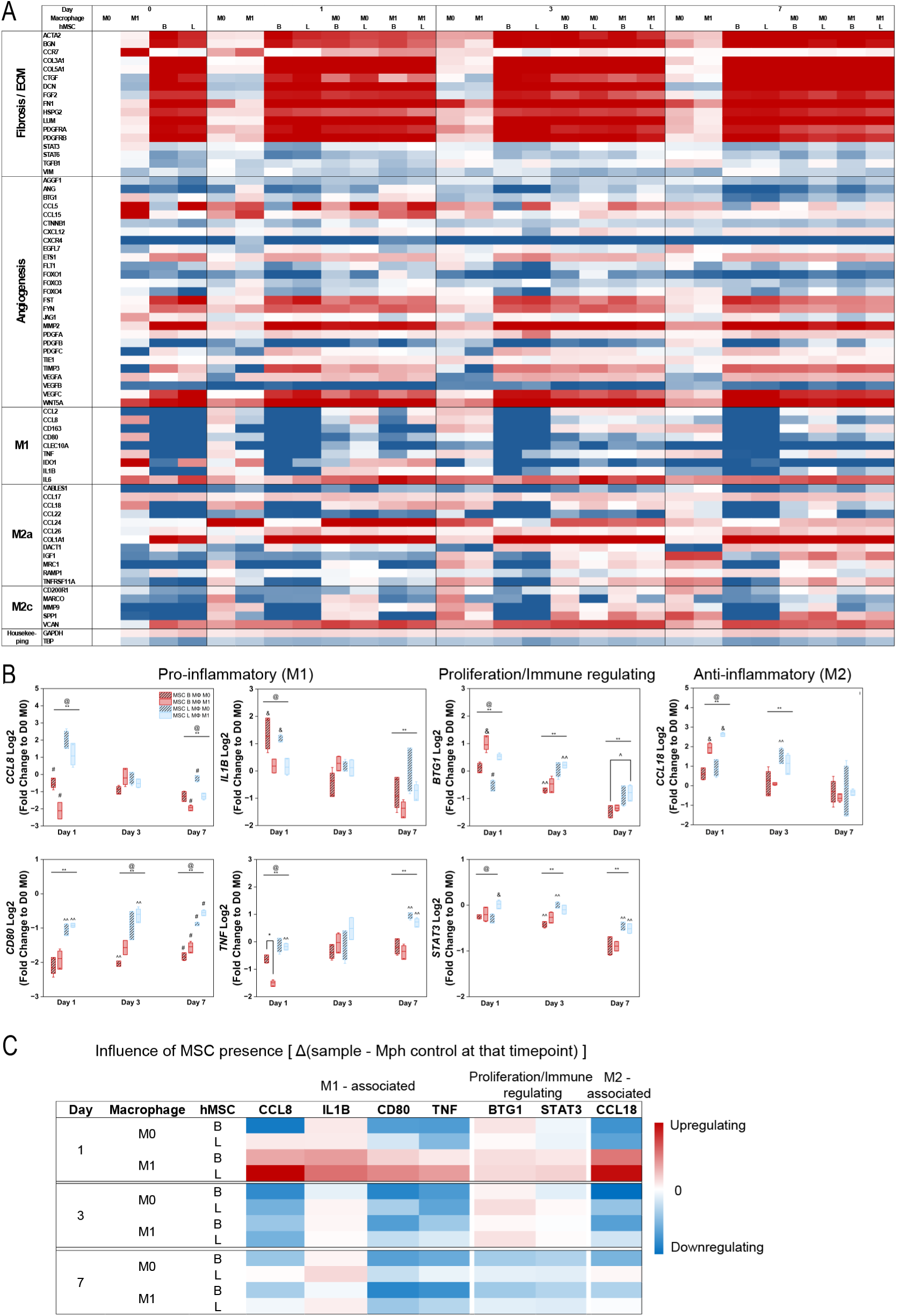
Patterns of macrophage gene expression as a function of inflammatory status and MSC co-culture. A custom NanoString code set was used to quantify macrophage gene expression in single culture controls and as a function of MSC co-culture conditions (basal vs. licensed). (A) All genes evaluated on a custom NanoString panel displayed as a fold change compared to M0 macrophage gene expression prior to seeding (n=3). A subset of genes was identified in the macrophage panel that were identified as macrophage only genes and where the co-cultures displayed > or < than 1 standard deviation from the single culture macrophage controls. (B) Co-culture gene expression displayed as a fold change compared to M0 macrophage gene expression prior to seeding (n=3). * denotes significance (p < 0.05) between indicated groups of same treatment (basal or licensed), ^ denotes significance (p < 0.05) between indicated groups of different treatment (basal or licensed), ** denotes overall significance (p < 0.05) between MSC treatments at the indicated timepoint, @ denotes overall significance (p < 0.05) between macrophage polarizations at the indicated timepoint, ^^ denotes significance (p < 0.05) between indicated group and all other groups of other MSC treatment, & denotes significance (p < 0.05) between indicated group and all other groups of other macrophage polarization, and # denotes significance (p < 0.05) of indicated group and all other groups at that timepoint. (C) The effect of MSCs co-cultures quantified as the difference of co-culture sample from macrophage-only controls at each timepoint. A red color indicates MSC co-culture enhances macrophage gene expression compared to a macrophage-only control. A blue color indicates MSC co-culture decreases macrophage gene expression compared to a macrophage-only control.

**Figure 5:**
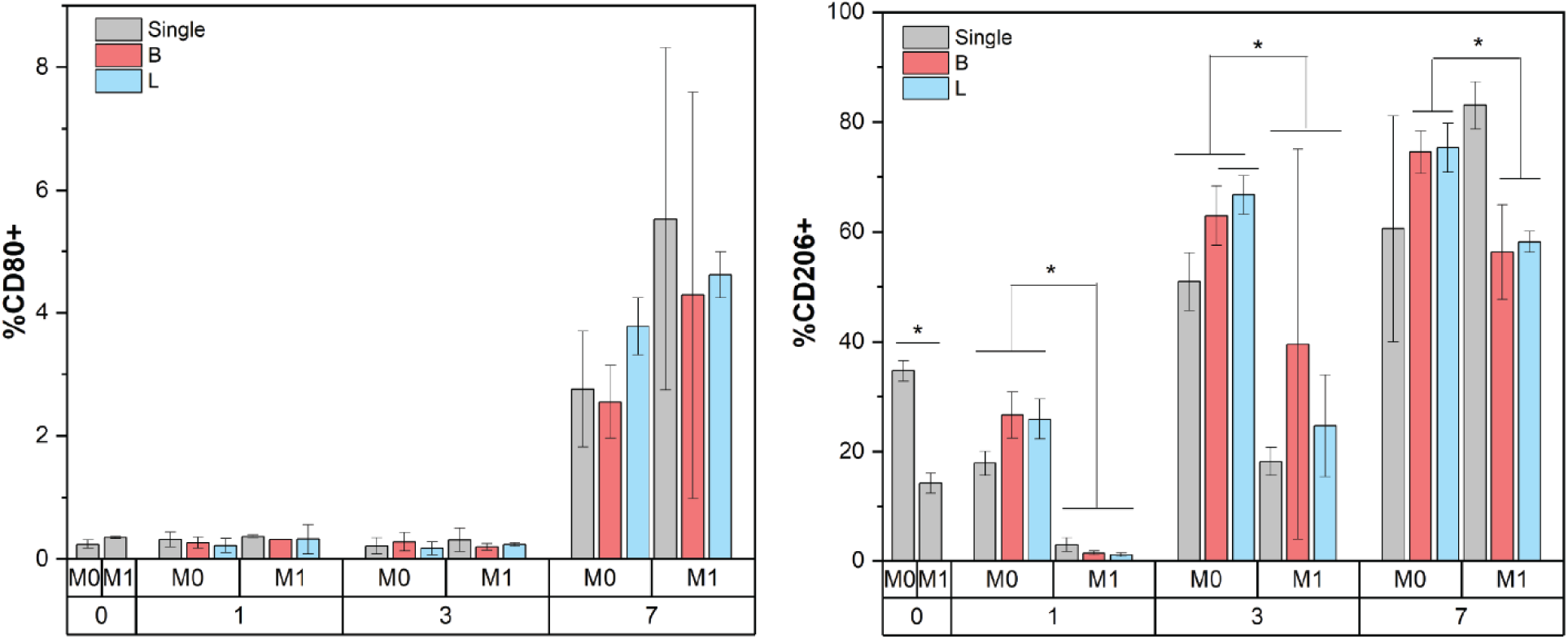
Macrophage surface marker expression was measured via flow cytometry. Two surface markers, CD80+ being an M1 indicator and CD206+ an M2 indicator, were chosen. Single cultures of M0 or M1 macrophages were conducted on mineralized collagen scaffolds and compared to co-cultures with basal or licensed MSCs (n=3). * indicates significance at p < 0.05 between M0 and M1 macrophages for indicated groups at the indicated timepoint.

The initial macrophage polarization state also significantly influenced subsequent macrophage activity within the scaffolds. Significant differences in gene expression were observed as early as day 1, where initially M0 macrophages expressed significantly higher *CCL8*, a pro-inflammatory factor which is known to induce M2-like macrophage polarization[38] and *IL1B*, which is known to be expressed in M1-like macrophages[39]. Cultures starting with M1 macrophages expressed significantly higher *BTG1*, which regulates cell growth, and *CCL18*, which is considered an M2-like gene[40, 41]. This suggests that while co-culture of M0 macrophages with MSCs on scaffolds enhances a pro-inflammatory phenotype, co-culture of M1 macrophages with MSCs on scaffolds can accelerate M2 polarization. Together, these findings suggest that in mineralized collagen scaffolds, recruitment of MSCs will temporarily polarize infiltrating pro-inflammatory macrophages to an anti-inflammatory (pro-regenerative) phenotype.

### 3.4 Macrophages display enhanced M2-like surface marker expression on mineralized collagen scaffolds with time

We then examined shifts in macrophage phenotype on scaffolds as a function of initial inflammatory stance as well as MSC co-culture via flow cytometry analysis of the M1 marker CD80[42] and the M2 marker CD206[43]. Cultures of macrophages seeded as M0 on the scaffolds displayed significantly greater expression of CD206 compared to macrophages seeded as M1, on days 1 and 3 **(Figure 5)**. Macrophages seeded as M0 co-cultured with MSCs also displayed significantly higher CD206 expression compared to macrophages seeded as M1 at all timepoints, regardless of MSC licensing. Interestingly, expression levels of CD206 and CD80 increased with culture time, reaching a maximum at day 7, with 50-80% of macrophages being CD206+ and 3-6% of macrophages being CD80+. No significant change in macrophage surface marker expression was observed as a function of MSC licensing status.

## 4. Discussion

The bone regeneration microenvironment is multicellular and undergoes significant change with time across the inflammatory phase after injury. Strategies that can shape this process may offer significant advantages for bone regeneration and remodeling. Tissue engineering and regenerative medicine approaches have traditionally sought to develop biomaterials that guide stem cell differentiation toward tissue-specific cells to enhance repair[44]. Here, we explored an alternative strategy leveraging the potential role MSCs play in multicellular interactions essential for shaping an immunomodulatory response. A key cell type in the inflammatory phase is the macrophage, which can take on a gradient of phenotypes largely categorized as either pro-inflammatory (M1) or pro-healing (M2)[45–47], although M2 macrophages are also associated with fibrosis[48–51]. MSCs have been shown to display an immunomodulatory response when licensed by inflammatory cytokines such as interferon-γ (IFN-γ) and tumor necrosis factor (TNF)[16, 20]. We previously showed the THP-1 macrophage secretome can educate MSCs towards a more immunomodulatory phenotype and enhance MSC osteogenesis regardless of MSC licensing state[22]. However, THP-1 derived macrophages display phenotypic differences compared to primary macrophages[52, 53]. In this study we used a well-characterized 3D mineralized collagen bone-mimetic biomaterial to investigate crosstalk between MSCs, initially in either an initially basal or licensed state, and primary human macrophages, initially in an unactivated (M0) or pro-inflammatory (M1) phenotype.

We co-cultured MSCs (basal vs. licensed) and macrophages (M0 vs. M1) on mineralized collagen scaffolds over 7 days and evaluated short-term MSC osteogenic and immunomodulatory responses as well as macrophage phenotypic expression. Our choice of culture time was informed both by the timeframe for MSC activation in our scaffolds as well as the reality of the wound environment immediately after scaffold implantation, which is known to strongly influence implant-defect integration[10, 21, 54]. Under normal healing conditions in the *in vivo* wound microenvironment, macrophages are known to start polarizing and gradually shift to reparative phenotypes (M2)[55]. In the scaffolds, basal MSCs showed increased secretion of osteoprotegerin (OPG), a key osteoclast inhibitor indicative of an osteogenic response, compared to licensed MSCs, and the presence of M1 macrophages enhanced this effect. This observation aligns with trends observed in our previous THP-1 – MSC co-culture experiments[22] and is consistent with previous observations that the mineralized collagen scaffold promotes rapid bone regeneration *in vivo*[19, 56]. We also evaluated the immunomodulatory capacity of MSCs in MSC-macrophage co-cultures. PGE_2_ secreted by licensed MSCs directly influences the activation, proliferation, differentiation and function of immune cells[17] and has been tied to increased IL-10 production by macrophages, indicative of an anti-inflammatory phenotype[57–61]. Our data show that groups containing licensed MSCs displayed enhanced PGE_2_ release on day 1 of culture (**Figure 2**). Interestingly, when compared to our prior co-cultures of MSCs with the THP-1 cell line, we observe that co-culture of primary macrophages with MSCs results in an order of magnitude greater PGE_2_ secretion. Another primary driver of MSC immunomodulatory effects is IL-6[17], a molecule with both pro- and anti-inflammatory effects and for which MSCs can be both a source and a target[62]. We observed a clear M1 macrophage-driven effect of IL-6 production in MSC-macrophage co-cultures, with greatest secretion in macrophage cultures with basal MSCs. Again, we observed a ten-fold increase in IL-6 expression in primary macrophage co-cultures as compared to our prior co-cultures using the THP-1 macrophage cell line[22]. This further cements the importance of evaluating macrophage response in regenerative biomaterials using primary macrophages, as common macrophage cell lines may not adequately capture the magnitude of responses. Taken together, this data shows that MSC licensing primarily guides the osteogenic potential of MSC-laden regenerative biomaterials, with basal MSCs showing greater pro-regenerative potency than licensed MSCs. However, and notably, inclusion of pro-inflammatory (M1-like) macrophages with MSCs can further increase secretion of immunomodulatory mediators such as IL-6.

A critical finding from this project is that macrophage co-culture upregulates basal MSC osteogenic and angiogenic gene expression by day 7 of culture compared to the MSC-only controls. We also evaluated macrophage gene expression and determined that while M1-associated genes were upregulated on the first day of culture in the scaffold, they subsequently showed a downregulation across the remainder of the culture period. Most notably, the inclusion of MSCs enhanced early M1-associated gene expression compared to macrophage-only culture controls, with the greatest effect seen for macrophages in co-culture with licensed MSCs. Although others have reported MSC-mediated macrophage polarization towards an M2 phenotype, these temporal dynamics in 3D microenvironments have not been reported previously. Evaluation of surface marker expression revealed increased M2-like macrophage (CD206) expression with culture time suggesting a time-dependent polarization pattern for macrophages in the scaffold, with a rapid transition towards an M2-like phenotype for M0 seeded groups and a slower transition for M1 seeded groups over a week in culture. Taken together, our findings suggest that a biomaterial platform designed to rapidly induce pro-inflammatory macrophage polarization may be highly beneficial to subsequently stimulate downstream MSC-mediated immunomodulatory and osteogenic activity that will in turn aid in macrophage polarization towards an anti-inflammatory phenotype and further promote regenerative healing.

Recent efforts have explored MSC -macrophage interactions in 3D biomaterials; however, these are hydrogel-based which fail to mimic the bone microenvironment[63]. This project offers a 2×2 experimental framework to functionally evaluate biomaterial performance through collective interrogation of inflammatory (licensed MSC, M1-like macrophage) and neutral (basal MSC, M0-like macrophage) multicellular interactions. We used a biomimetic scaffold with demonstrated capacity to induce craniofacial bone regeneration *in vivo*. Compositional[21, 29, 64], structural[20], and mechanical[31, 65] modifications to the mineralized collagen scaffold have been extensively described by our lab in the context of shaping patterns of MSC osteogenesis. However, a greater understanding of how the biomaterial properties directly or indirectly (via paracrine signals from other cells like osteoprogenitors) influence macrophage polarization represents a significant opportunity to improve regenerative potential. Future studies will focus on evaluating macrophage polarization as a function of material composition and structure to elucidate material-cell interactions. This may be particularly significant in the context of varying the proteoglycan content of these scaffolds, which may directly act to shape macrophage polarization but may also act indirectly via charge-mediated interactions that alter the sequestration and bioavailability of biomolecular signals[21, 66, 67] that contribute to immunomodulatory activity. Similarly, modifications to the scaffold mineral content such as recent incorporation of zinc ions that alter the morphology of brushite mineral deposits from plate-like to needle-like[68], may significantly alter macrophage polarization, given prior work showing the importance of alignment on macrophage phenotype[30, 69]. Further studies are also needed to evaluate patient-to-patient variability between the single and co-culture conditions of MSCs and macrophages. We have recently published a thorough study evaluating donor variability in MSC osteogenic response within these same mineralized collagen scaffolds[70], so a similar study evaluating donor variability in regards to primary macrophages would be impactful to the field. Importantly, the use of a standardized 3D biomaterial that replicates key aspects of the native tissue microenvironment is likely to provide significantly greater insight than more conventional 2D and conditioned media experiments that have been performed to date.

## 5. Conclusions

MSC interactions with macrophages have garnered increasing attention in the search for strategies to modulate the wound microenvironment and shape the quality and kinetics of bone repair. Here we explored the influence of direct co-culture of hMSCs in an initially basal or licensed state with primary human macrophages in a neutral or pro-inflammatory state within a mineralized collagen biomaterial. We show MSC osteogenic and immunomodulatory response as well as macrophage polarization kinetics are strongly influenced by both initial MSC or macrophage state but also by reciprocal interactions between these populations within the collagen biomaterial in as fast as 7 days. Notably, initial MSC licensing status is a significant factor shaping both MSC and macrophage response. Initially, basal MSCs display a stronger osteogenic response while inflammatory licensed MSCs display a strong immunomodulatory response. However, with time in culture, the presence of macrophages is responsible for shaping significant shifts in regenerative gene expression. Further, while macrophages cultured on mineralized collagen scaffolds initially display signs of adopting a pro-inflammatory phenotype, co-culture with MSCs further upregulates this, with the greatest effects for MSCs already licensed by inflammatory stimuli. Interestingly, macrophage pro-inflammatory status dampens with time, suggesting a phenotypic transition towards a more reparative macrophage phenotype within the mineralized collagen scaffold. This work expands our understanding of MSC-macrophage interactions and illuminates the opportunity to exploit this relationship for improved biomaterial design.

## Acknowledgements

The authors would like to acknowledge the following institutes for access to their facilities and services: the School of Chemical Sciences Microanalysis Laboratory, the Carl R. Woese Institute for Genomic Biology, the Tumor Engineering and Phenotyping Shared Resource (TEP) at the Cancer Center at Illinois, and the Beckman Institute for Advanced Science and Technology, located at the University of Illinois. Research reported in this publication was supported by the National Institute of Dental and Craniofacial Research of the National Institutes of Health under Award Number R21 DE026582 and R01 DE030491 (BACH), National Institute of Arthritis and Musculoskeletal and Skin Diseases under Award Number R01 AR077858 (BACH), and National Heart, Lung, and Blood Institute R01 HL130037 (KLS). We are also grateful for funds provided by the NSF Graduate Research Fellowship (DGE-174604 to VK; DGE-1144245 to AST) and the Chemistry-Biology Interface Research Training Program at the University of Illinois (T32 GM070421, VK). Additional support was provided by the Carl R. Woese Institute for Genomic Biology and the Chemical and Biomolecular Engineering Dept. at the University of Illinois at Urbana-Champaign. The interpretations and conclusions presented are those of the authors and are not necessarily endorsed by the National Institutes of Health or the National Science Foundation.

## Contributions (CRediT: Contributor Roles Taxonomy)

**Vasiliki Kolliopoulos:** Conceptualization, Data curation, Formal Analysis, Visualization, Investigation, Methodology, Writing – original draft, Writing – review & editing. **Maxwell Polanek:** Methodology, Formal Analysis, Visualization, Investigation, Writing – original draft, Writing – review & editing. **Hashni E. Vidana Gamage:** Investigation, Formal Analysis. **Melisande Wong Yan Ling:** Investigation. **Aleczandria Tiffany:** Investigation. **Erik Nelson:** Supervision, Methodology, Writing – review & editing. **Kara Spiller:** Supervision, Methodology, Writing – review & editing. **Brendan Harley:** Conceptualization, Resources, Project administration, Funding acquisition, Supervision, Writing – review & editing.

## Data Availability

The raw data and processed data required to reproduce these findings are available upon request to Brendan Harley.

## Disclosure

The authors have no conflicts of interest to report.

## Notes

### Competing Interest Statement

The authors have declared no competing interest.

