## Supplemental Info for "Reciprocal macrophage-MSC crosstalk drives immunomodulatory and regenerative phenotypes in a mineralized collagen scaffold"

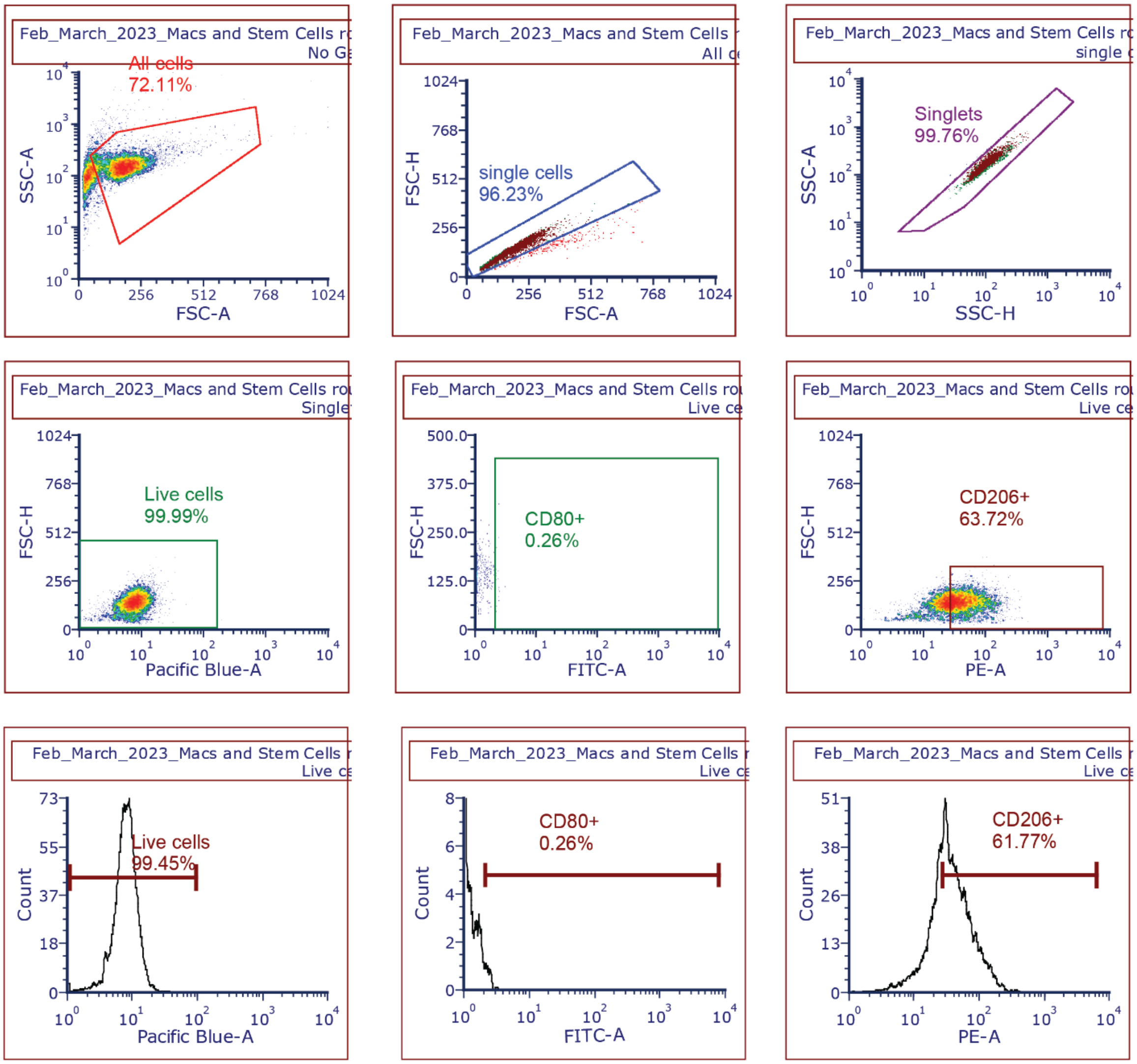


**Supp. Figure 1**: Macrophage surface marker expression was measured via flow cytometry. Two surface markers, CD80+ being an M1 indicator, and CD206+ an M2 indicator were chosen along with a live/dead stain. A representative gating strategy used for all samples is depicted here.

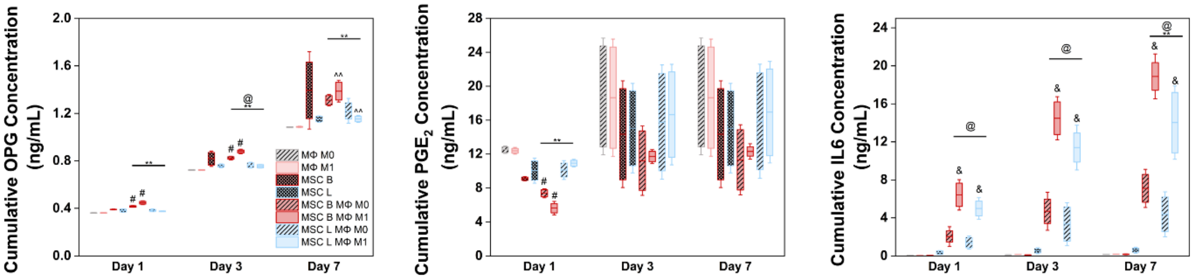


**Supp. Figure 2**: Mesenchymal stem cell (MSC) and macrophage (MΦ ) secreted factors measured via ELISA. The following statistical symbols indicate significance at $p<0.05$: ^^: Indicated group is significantly different from both the groups of opposite MSC treatment, &: Indicated group is significantly different from both the groups of opposite Macrophage polarization, **: Overall difference between the behavior of licensed and basal MSCs at this time point, @: Overall difference between the behavior of M1 and M0 macrophages at this time point, and #: Indicated group is different from all other co-cultures at that time point.

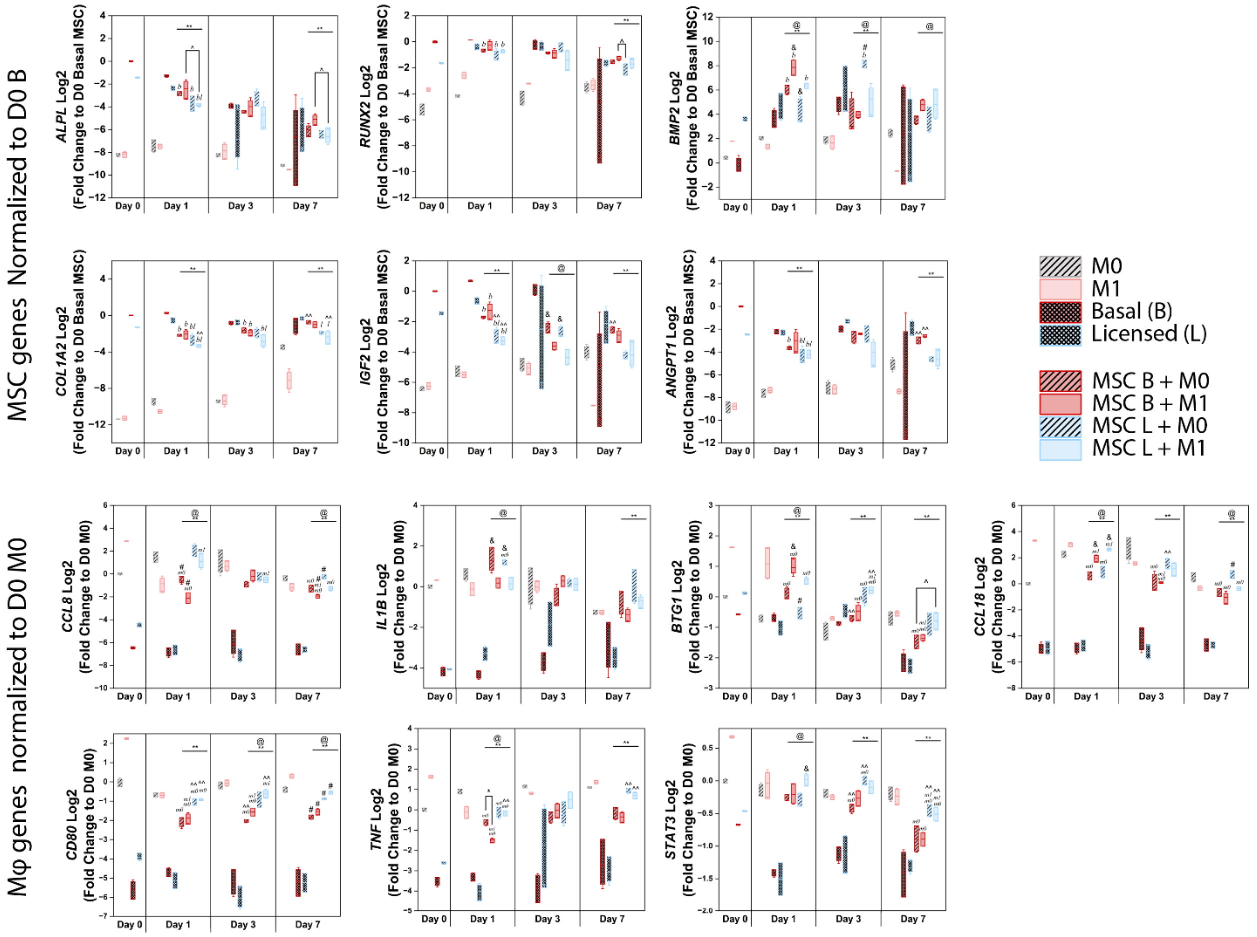


**Supp. Figure 3**: Mesenchymal stem cell (MSC) and macrophage (MΦ ) N anoS tring gene expression depicted with all single and co-cultures. MSC gene expression is normalized to the day 0 MSC basal single culture expression while MΦ gene expression is normalized to the day 0 M0 single culture expression and each are presented as a fold change. The following statistical symbols indicate significance at $p<0.05$: *: indicated groups are significantly different (within same MSC treatment), ^: indicated groups are significantly different (between different MSC treatments), ^^: Indicated group is significantly different from both the groups of opposite MSC treatment, &: Indicated group is significantly different from both the groups of opposite Macrophage polarization, **: Overall difference between the behavior of licensed and basal MSCs at this time point, @: Overall difference between the behavior of M1 and M0 macrophages at this time point, #: Indicated group is different from all other co-cultures at that time point, *m0*: Indicated group is different from the M0 macrophage single culture control, *m1*: Indicated group is different from the M1 macrophage single culture control, *b*: Indicated group is different from the basal MSC single culture control, *l*: Indicated group is different from the licensed MSC single culture control.

**Supp. Table 1**: MSC NanoString panel gene list.

| Customer Name | HUGO Gene | Probe NSID |
|---|---|---|
| ALPL | ALPL | NM_000478.3:2065 |
| ANGPT1 | ANGPT1 | NM_001146.3:2080 |
| BGLAP | BGLAP | NM_199173.4:44 |
| BMP2 | BMP2 | NM_001200.2:1515 |
| BMP7 | BMP7 | NM_001719.1:525 |
| CCL2 | CCL2 | NM_002982.3:123 |
| CCL7 | CCL7 | NM_006273.2:120 |
| COL1A2 | COL1A2 | NM_000089.3:2635 |
| MCSF | CSF1 | NM_000757.4:823 |
| IL8 | CXCL8 | NM_000584.2:25 |
| FGFR2 | FGFR2 | NM_000141.4:2204 |
| GAPDH | GAPDH | NM_001256799.1:386 |
| GUSB | GUSB | NM_000181.3:1899 |
| HGF | HGF | NM_000601.4:550 |
| IDO1 | IDO1 | NM_002164.5:369 |
| IGF2 | IGF2 | NM_000612.4:765 |
| IHH | IHH | NM_002181.2:1693 |
| IL10 | IL10 | NM_000572.2:622 |
| IL1RN | IL1RN | NM_000577.3:480 |
| IL6 | IL6 | NM_000600.3:364 |
| GAL9 | LGALS9 | NM_002308.3:359 |
| MMP9 | MMP9 | NM_004994.2:1530 |
| OAZ1 | OAZ1 | NM_004152.2:313 |
| PTGS2 | PTGS2 | NM_000963.1:495 |
| RUNX2 | RUNX2 | NM_004348.3:1850 |
| SEMA3A | SEMA3A | NM_006080.1:585 |
| SMAD5 | SMAD5 | NM_005903.5:1044 |
| SOX9 | SOX9 | NM_000346.2:2135 |
| SP7 | SP7 | NM_001173467.1:1510 |
| OPN | SPP1 | NM_000582.2:760 |
| TSG6 | TNFAIP6 | NM_007115.2:250 |
| TNFRSF11A | TNFRSF11A | NM_003839.3:226 |
| TNFRSF11B | TNFRSF11B | NM_002546.2:1075 |
| TNFSF11 | TNFSF11 | NM_003701.2:490 |
| VEGFA | VEGFA | NM_001025366.1:1325 |
| VEGFB | VEGFB | NM_003377.3:687 |
| WNT16 | WNT16 | NM_057168.1:1621 |
| WNT5a | WNT5A | NM_003392.3:475 |

**Supp. Table 2**:MΦ NanoString panel gene list.

| Customer Name | HUGO Gene | Probe NSID |
|---|---|---|
| ACTA2 | ACTA2 | NM_001613.1:645 |
| AGGF1 | AGGF1 | NM_018046.3:35 |
| ANG | ANG | NM_001145.4:949 |
| BGN | BGN | NM_001711.3:1935 |
| BTG1 | BTG1 | NM_001731.2:775 |
| CABLES1 | CABLES1 | NM_001100619.2:2560 |
| CCL15 | CCL15 | NM_032965.4:869 |
| CCL17 | CCL17 | NM_002987.2:229 |
| CCL18 | CCL18 | NM_002988.2:585 |
| CCL2 | CCL2 | NM_002982.3:123 |
| CCL22 | CCL22 | NM_002990.3:797 |
| CCL24 | CCL24 | NM_002991.2:18 |
| CCL26 | CCL26 | NM_006072.4:184 |
| CCL5 | CCL5 | NM_002985.2:280 |
| CCL8 | CCL8 | NM_005623.2:689 |
| CTGF | CCN2 | NM_001901.2:1100 |
| CCR7 | CCR7 | NM_001838.2:1610 |
| CD163 | CD163 | NM_004244.4:1630 |
| CD200R1 | CD200R1 | NM_138806.3:142 |
| CD80 | CD80 | NM_005191.3:674 |
| CLEC10A | CLEC10A | NM_182906.2:430 |
| COL1A1 | COL1A1 | NM_000088.3:5210 |
| COL3A1 | COL3A1 | NM_000090.3:180 |
| COL5A1 | COL5A1 | NM_000093.3:872 |
| CTNNB1 | CTNNB1 | NM_001098210.1:1815 |
| CXCL12 | CXCL12 | NM_199168.3:69 |
| CXCR4 | CXCR4 | NM_003467.2:1335 |
| DACT1 | DACT1 | NM_001079520.1:3350 |
| DCN | DCN | NM_001920.3:420 |
| EGFL7 | EGFL7 | NM_016215.3:1252 |
| ETS1 | ETS1 | NM_005238.3:1305 |
| FGF2 | FGF2 | NM_002006.4:620 |
| FLT1 | FLT1 | NM_002019.4:530 |
| FN1 | FN1 | NM_212482.1:1776 |
| FOXO1 | FOXO1 | NM_002015.3:1526 |
| FOXO3 | FOXO3 | NM_001455.2:1860 |
| FOXO4 | FOXO4 | NM_001170931.1:1121 |
| FST | FST | NM_006350.2:575 |
| FYN | FYN | NM_002037.3:765 |

| | | |
|---|---|---|
| GAPDH | GAPDH | NM_001256799.1:386 |
| HSPG2 | HSPG2 | NM_005529.5:2715 |
| IDO1 | IDO1 | NM_002164.5:369 |
| IGF1 | IGF1 | NM_000618.3:491 |
| IL1B | IL1B | NM_000576.2:840 |
| IL6 | IL6 | NM_000600.3:364 |
| JAG1 | JAG1 | NM_000214.2:915 |
| LUM | LUM | NM_002345.3:1285 |
| MARCO | MARCO | NM_006770.3:61 |
| MMP2 | MMP2 | NM_004530.2:2360 |
| MMP9 | MMP9 | NM_004994.2:1530 |
| MRC1 | MRC1 | NM_002438.2:525 |
| PDGFA | PDGFA | NM_002607.5:2460 |
| PDGFB | PDGFB | NM_033016.2:1480 |
| PDGFC | PDGFC | NM_016205.2:2596 |
| PDGFRA | PDGFRA | NM_006206.3:1925 |
| PDGFRB | PDGFRB | NM_002609.3:265 |
| RAMP1 | RAMP1 | NM_005855.2:200 |
| SPP1 | SPP1 | NM_000582.2:760 |
| STAT3 | STAT3 | NM_003150.3:2060 |
| STAT6 | STAT6 | NM_003153.3:2030 |
| TBP | TBP | NM_001172085.1:587 |
| TGFB1 | TGFB1 | NM_000660.3:1260 |
| TIE1 | TIE1 | NM_005424.2:2610 |
| TIMP3 | TIMP3 | NM_000362.4:1640 |
| TNF | TNF | NM_000594.2:1010 |
| TNFRSF11A | TNFRSF11A | NM_003839.3:226 |
| VCAN | VCAN | NM_004385.3:9915 |
| VEGFA | VEGFA | NM_001025366.1:1325 |
| VEGFB | VEGFB | NM_003377.3:687 |
| VEGFC | VEGFC | NM_005429.2:565 |
| VIM | VIM | NM_003380.2:694 |
| WNT5A | WNT5A | NM_003392.3:475 |

**Supp. Table 3**: Macrophage surface marker expression was measured via flow cytometry. Two surface markers, CD80+ being an M1 indicator, and CD206+ an M2 indicator were chosen along with a live dead viability stain and Fc blocker.

| Target | Species | Conjugate | Source | Dilution |
|---|---|---|---|---|
| CD80 | Human | FITC | eBioscience | 100x |
| CD206 | Human | PE | eBioscience | 100x |
| LIVE/DEAD Fixable | N/A | Violet | ThermoFisher | 1000x |
| Fc receptor binding inhibitor | Human | Unconjugated | eBioscience | |
